# Diurnal changes in perineuronal nets and parvalbumin neurons in the rat medial prefrontal cortex

**DOI:** 10.1101/2020.10.25.354134

**Authors:** John H. Harkness, Angela E. Gonzalez, Priyanka N. Bushana, Emily T. Jorgensen, Deborah M. Hegarty, Ariel A. Di Nardo, Alain Prochiantz, Jonathan P. Wisor, Sue A. Aicher, Travis E. Brown, Barbara A. Sorg

## Abstract

Perineuronal nets (PNNs) surrounding fast-spiking, parvalbumin (PV) inhibitory interneurons are vital for providing excitatory:inhibitory balance within cortical circuits, and this balance is impaired in disorders such as schizophrenia, autism spectrum disorder, and substance use disorders. These disorders are also associated with altered diurnal rhythms, yet few studies have examined the diurnal rhythms of PNNs or PV cells. We measured the intensity and number of PV cells and PNNs labeled with *Wisteria floribunda* agglutinin (WFA) in the rat prelimbic medial prefrontal cortex (mPFC) at Zeitgeber times (ZT) ZT0, 6, 12, and 18. We also measured the oxidative stress marker 8-oxo-deoxyguanosine (8-oxo-dG). Relative to ZT0, the intensities of PNN and PV staining were increased in the dark (active) phase compared with the light (inactive) phase. The intensity of 8-oxo-dG was decreased from ZT0 at all time points (ZT6,12,18), in both PV cells and non-PV cells. To examine corresponding changes in inhibitory and excitatory inputs, we measured GAD 65/67 and vGlut1 puncta apposed to PV cells with and without PNNs. Relative to ZT6, there were more excitatory puncta on PV cells surrounded by PNNs at ZT18, but no changes in PV cells devoid of PNNs. No changes in inhibitory puncta were observed. Whole-cell slice recordings in fast-spiking (PV) cells with PNNs showed an increased ratio of α-amino-3-hydroxy-5-methyl-4-isoxazolepropionic acid receptor:N-methyl-D-aspartate receptor (AMPA:NMDA) at ZT18 *vs*. ZT6. The number of PV cells and co-labeled PV/PNN cells containing the transcription factor orthodenticle homeobox 2 (OTX2), which maintains PNNs, showed a strong trend toward an increase from ZT6 to ZT18. These diurnal fluctuations in PNNs and PV cells are expected to alter cortical excitatory:inhibitory balance and provide new insights into treatment approaches for diseases impacted by imbalances in sleep and circadian rhythms.

## INTRODUCTION

Perineuronal nets (PNNs) are specialized extracellular matrix structures that surround specific neurons in the brain and spinal cord [1], appear during critical periods of development [2–5], and restrict plasticity during adulthood [6]. PNNs surround mainly parvalbumin (PV)-containing, fast-spiking GABAergic interneurons in several brain regions [7], including in the medial prefrontal cortex (mPFC) [8]. The prelimbic region of the mPFC is associated with working memory and cognition [9, 10]. PV neurons profoundly inhibit the network of surrounding neurons via their elaborate contacts with local pyramidal neurons [11], thereby regulating plasticity associated with learning, decision making, attention, cognitive flexibility, and working memory [12–15]. PV cells therefore contribute to essential excitatory:inhibitory balance for normal functioning of the mPFC [12] and are critical for generating gamma oscillations [16–18] believed to mediate normal working memory function and cognitive flexibility [13, 19].

Perineuronal nets and PV neurons are important for normal learning and memory processes, and their dysfunction appears to contribute to a wide range of brain diseases/disorders, including schizophrenia, bipolar disorder, Alzheimer’s disease, autism spectrum disorder, epilepsy, and disorders associated with drugs of abuse and fear [20–30]. Several of these disorders and cognitive impairments are also associated with circadian and/or sleep disturbances: most notably schizophrenia [31–33], autism spectrum disorder [34], epilepsy [35], drug use disorders [36–38], and neurodegenerative diseases such as Alzheimer’s disease [39, 40]. These disorders are all accompanied by poor working memory function and cognitive flexibility [41–44], functions that depend on optimal excitatory:inhibitory balance in the mPFC. The dynamics of cortical excitability are dependent on sleep homeostatic drive, which increases as the duration of prior wakefulness increases [45]. More recently, these sleep-related effects are responsive to key dynamics in circadian rhythms, including the amplitude of these rhythms [46, 47]. Thus, a fundamental understanding of how PNNs and their underlying PV interneurons contribute to plasticity in response to outside stimuli, including diurnal physiological changes, is expected to provide new avenues for understanding and regulating excitatory:inhibitory balance in the mPFC.

We previously showed that sleep disruption in rats increased the intensity of the PNN marker, *Wisteria floribunda* agglutinin (WFA), PV, and the oxidative stress marker 8-oxo-2’-deoxyguanosine (8-oxo-dG) [48]. However, the magnitude of WFA changes were small, and we examined only one time of day when comparing sleep-disrupted rats with undisrupted controls. It is possible that these changes in marker intensity were masked by time-of-day changes created by diurnal fluctuations. Pantazopolous *et al.* [49] recently reported that the number of PNNs in mice fluctuated diurnally in several brain regions, including the prelimbic mPFC, and that this same pattern persisted when mice were maintained under constant darkness, indicating a circadian influence. Here we tested the hypotheses that both PNN and PV intensity fluctuate diurnally. These fluctuations may reflect increased activity levels of rats during their active periods while facilitating greater synaptic plasticity during sleep. PNN and PV intensity were quantified in the prelimbic mPFC at four times of day (ZT0, ZT6, ZT12, ZT18). In addition, we measured the intensity of 8-oxo-dG as an indicator of oxidative stress. We also hypothesized that glutamatergic signaling in WFA+/PV+ neurons would cycle between the light (ZT6) *vs*. the dark cycle (ZT18) and therefore we measured GAD65, GAD67, vGLUT1, and the volume of these neurons in the prelimbic mPFC at the same time points. Additionally, we measured the AMPA:NMDA ratio in fast-spiking/WFA+ neurons at ZT6 and ZT18 to determine if changes in excitatory inputs were reflected in changes in neuronal activity. Finally, at ZT6 and ZT18, we also assessed the number of cells containing orthodenticle homeobox 2 (OTX2), a homeoprotein transcription factor required for normal development of the CNS [50, 51] that is imported by PNN-enwrapped PV cells [52–55]. Our findings collectively indicate that several parameters of PV cells fluctuate diurnally, and these fluctuations have implications for altered excitatory:inhibitory balance in brain disorders with disrupted circadian rhythms and sleep.

## METHODS

### Animals

A total of 48 rats were used for the experiments. Male Sprague Dawley rats weighing 260-280 g were obtained from Envigo (Livermore, CA) and individually housed under LD12:12 conditions. Rats were maintained between 18° and 22°C and a relative humidity of 55%. Rats were allowed to acclimate to the light cycle for two weeks prior to euthanizing at four different times of day (see below). Protocols were approved by the Washington State University Institutional Animal Care and Use Committee (Protocol #07432). All efforts were made to reduce animal suffering.

### Experimental design

Rats were euthanized every 6 hours over four different time points, including Zeitgeber time (ZT)0, at lights-on, and ZT6, ZT12, and ZT18 (N = ZT0 (10), ZT6 (8), ZT12 (7), ZT18 (8)). Rats were removed from the colony one at a time and within 30 sec to 1 min after removal, they were given an overdose of pentobarbital (50 mg/kg under brief, 5% isoflurane exposure) at the designated time point.

### Immunohistochemistry

Upon reaching unresponsiveness from pentobarbital overdose, rats were perfused transcardially with 150 mL 0.1M phosphate-buffered saline (PBS) at a rate of 300 mL/min. Perfusate was switched to 4% paraformaldehyde and rats were perfused at the same rate for 250 mL/rat. Brains were removed, immersed in 20 mL 4% paraformaldehyde overnight, and then immersed in 20% sucrose in PBS solution and refrigerated. After two days in sucrose solution (when the brains sank), brains were flash-frozen with powdered dry ice and stored at −80°C until sectioned.

Coronal sections containing the prelimbic PFC from +3.2 to +4.2 mm from bregma [56] were collected on a freezing microtome at 30 μm for a 1:8 section series [57]. Triple-staining was performed by first washing a single series of free-floating sections three times for 5 min in PBS. Tissue was then treated with 50% ethanol for 30 min. Sections were washed in PBS three times for 5 min each before being placed in a blocking solution containing 3% normal goat serum (Vector Laboratories) for 1 hr. Subsequently, tissue was co-incubated with rabbit-anti-PV (ThermoFisher Scientific, Cat# PA1-933, RRID: AB_2173898,1:1000) and mouse-anti-8-oxo-dG (EMD Millipore, Cat# 4354-MC-050, RRID: AB_2876794,1:350) and 2% normal goat serum at 4°C overnight. Tissue was then rinsed in PBS three times for 10 min each and incubated for 2 hr in secondary antibodies (goat anti-rabbit Alexa Fluor® 405 for the PV antibody, (abcam, Cat# ab175652, RRID AB_2687498, 1:500), goat anti-mouse Alexa Fluor® 594 for the 8-oxo-dG antibody (abcam, Cat# ab150120, RRID AB_2650601, 1:500), and 2% normal goat serum. Slices were washed in PBS three times for 10 min each and then incubated with the fluorescein isothiocyanate (FITC)-conjugated PNN marker, *Wisteria floribunda* agglutinin (WFA, Vector Laboratories; Cat# FL-1351, AB_2336875, 1:500), which is widely used to label PNNs [1], and 2% normal goat serum at 4°C overnight. After three 10 min washes in PBS, sections were mounted onto Superfrost Plus slides and allowed to dry overnight. After drying, ProLong Gold Antifade Mountant (ThermoFisher Scientific) was applied to the slides before coverslipping.

We also measured OTX2 staining in a subset of brain slices used to measure WFA and PV. Owing to constraints in the protocol, we did not make comparisons of cell intensity and instead analyzed only cell numbers. After mounting to Superfrost Plus slides and brief drying, tissue sections were pretreated with 0.5% Triton-X and PBS solution followed by 100 mM glycine. Slides were washed three times with PBS, then bathed in blocking solution containing 5% BSA (Sigma) and 0.5% Triton-X in PBS for 30 minutes. Tissue was co-incubated with rabbit-anti-PV (as above) and mouse-anti-OTX-2 (1:25, in-house, Prochiantz Laboratory) in 5% BSA, 0.5% Triton-X in PBS in a humidified chamber at 4°C overnight. Tissue was then rinsed in PBS for 5 seconds and washed three times in PBS for 10 min each. Slices were incubated for 2 hr in a secondary antibody cocktail (goat anti-rabbit Alexa Fluor® 405 for the PV antibody and goat anti-mouse Alexa Fluor® 594 for the OTX-2 antibody, 1:500) in 5% BSA, 0.5% Triton-X in PBS. Tissue was then washed in PBS for 5 seconds, and three times for 10 min each. Finally, tissue was incubated in FITC-WFA (as above) in 5% BSA, 0.5% Triton-X in PBS for 2 hr at room temperature, washed in PBS for 5 seconds, and then washed three times for 10 min each. After drying, ProLong Gold Antifade Mountant (ThermoFisher Scientific) was applied to the slides before coverslipping.

### Quantification of immunohistochemical images

Imaging for WFA, PV, 8-oxo-dG, and OTX2 was performed on a Leica SP8 laser scanning confocal microscope with an HCX PL apo CS, dry, 20x objective with 0.70 numerical aperture. 405, 488, and 594 nm lasers were used for excitation, and were detected by three photomultiplier tubes in the 400-450, 460-510, and 590-640 nm ranges, respectively. Calibration of the laser intensity, gain, offset, and pinhole settings were determined within the orbitofrontal cortex of a control animal, as this region most reliably expresses strong WFA staining. These settings were maintained for all images. Images were collected in z-stacks of 20 images each (step size 0.44 μm; containing the middle 8.45 μm of each brain section), encompassing the prelimbic PFC.

All images (1.194 pixels/μm; 428 × 428 µm) were compiled into summed images using ImageJ macro plug-in Pipsqueak AI™ (https://pipsqueak.ai) [58], scaled, and converted into 8-bit, grayscale, tiff files. Pipsqueak AI™ was run in “semi-automatic mode” to select ROIs to identify individual PV+ cells, PNNs, 8-oxo-dG, or OTX2-labeled cells, which were then verified by a trained experimenter who was blinded to the experimental conditions. The plug-in compiles this analysis to identify single-[59], double-labeled [60], and triple-labeled [48] neurons. Labeling was quantified bilaterally in the prelimbic mPFC using Pipsqueak AI software. Background threshold levels were set and applied to all images for comparison. When describing co-labeling of PNNs with another cell marker (e.g., PV, OTX2 double- or triple-labeling), we acknowledge that WFA labeling is not strictly co-labeled but instead located around the perimeter of the markers located inside the cell.

### Quantification of PV puncta and cell volume

Because we generally found the maximal differences in PV and WFA labeling intensity between ZT6 (nadir) and ZT18 (peak), we assessed the number of excitatory and inhibitory puncta on PV cells surrounded by PNNs at these two times of day. Immunohistochemistry and imaging were performed on brain slices from a subset of the animals used for WFA and PV intensity studies. Immunohistochemical methods were similar to those previously described [60–63]. Solutions were prepared in either 0.1 M phosphate buffer at pH 7.4 (PB) or 0.1 M Tris-buffered saline at pH 7.6 (TS). Tissue sections were first rinsed in PB, then incubated in 1% sodium borohydride in PB for 30 min to reduce background. After rinses in PB and TS, sections were incubated in 0.5% bovine serum albumin (BSA) in TS for 30 min and then placed in a primary antibody cocktail made in 0.1% BSA and 0.25% Triton X-100 in TS for two nights at 4°C. The primary antibody cocktail consisted of mouse anti-glutamic acid decarboxylase 65 (GAD65, abcam, Cat# ab26113, RRID: AB_448989, 1:500), mouse anti-glutamic acid decarboxylase 67 (GAD67, Millipore Sigma, Cat# MAB5406, RRID: AB_2278725, 1:1000), rabbit anti-PV (Novus Biologicals, Cat# NB120-11427, RRID:AB_791498, 1:1000), and guinea pig anti-vesicular glutamate transporter 1 (vGLUT1; EMD Millipore, Cat# AB5905, RRID: AB_2301751, 1:5000). After 40 h primary antibody incubation, tissue sections were rinsed in TS and then incubated with a cocktail of fluorescently-labeled secondary antibodies for 2 h, light-protected, at room temperature. The secondary antibody cocktail consisted of Alexa Fluor 488 donkey anti-mouse (ThermoFisher Scientific, Cat# A21202, RRID: AB_141607, 1:800) to label both GAD antibodies, Alexa Fluor 546 donkey anti-rabbit (ThermoFisher Scientific, Cat# A10040, RRID: AB_2534016, 1:800;); and Alexa Fluor 647 donkey anti-guinea pig (Jackson ImmunoResearch Laboratories, Cat# 706-605-148, RRID: AB_2340476, 1:800). Tissue sections were rinsed again in TS and then incubated in biotinylated WFA (Vector Laboratories, Cat# B-1355, RRID: AB_2336874, 1:50) for 2 h at room temperature. Following TS rinses, tissue sections were incubated for 3 h at room temperature in Alexa Fluor 405-conjugated streptavidin (ThermoFisher Scientific, Cat# S32351, 6.25 μg/ml). Finally, tissue sections were rinsed in TS followed by PB before being mounted with 0.05 M PB onto gelatin-coated slides to dry. Slides were coverslipped with Prolong Gold Antifade Mountant (ThermoFisher Scientific) and light-protected until imaging. Anatomical landmarks were used to determine representative caudal and rostral sections of prelimbic cortex that were within bregma +3.5 to +4.2 mm [56].

Confocal imaging was performed as described previously [60, 61]. Two high magnification images were taken at each rostral-caudal level of the prelimbic mPFC (2 images/level × 2 levels/animal = 4 images/animal). Images were captured on a Zeiss LSM 780 confocal microscope with a 63 × 1.4 NA Plan-Apochromat objective (Carl Zeiss MicroImaging, Thornwood, NY) using the single pass, multi-tracking format at a 1024 × 1024 pixel resolution. Optical sectioning produced Z-stacks bounded by the extent of fluorescent immunolabeling throughout the thickness of each section. Using Zen software (Carl Zeiss, RRID SCR_013672), PV neurons in each confocal stack were identified and assessed for the presence of a nucleus and whether the entire neuron was within the boundaries of the field of view; only these PV neurons were included in the analysis. The optical slice through the nucleus at which the ellipsoidal minor axis length of each PV neuron reached its maximum was determined. A Z-stack of that optical slice plus one optical slice above and one below was created resulting in a 1.15 μm Z-stack through the middle of each PV neuron; these subset Z-stacks were used for puncta apposition analysis.

Image analysis of GABAergic and glutamatergic appositions onto PV-labeled neurons was performed as described [61] using Imaris 9.0 software (BitPlane USA, Concord, MA, RRID: SCR_007370) on an offline workstation in the Advanced Light Microscopy Core at Oregon Health & Science University by an observer who did not know experimental conditions. For each PV neuron, the manual setting of the *Surfaces* segmentation tool was used to trace the outline of the PV neuron in all three optical slices and a surface was created. The volume of the PV cell rendered model was measured by Imaris. To limit our analyses to the area immediately surrounding each PV neuron, we used the *Distance Transform* function followed by the automated *Surfaces* segmentation tool to create another surface 1.5 μm away from the PV neuron surface that followed the unique contours of that PV neuron. The *Mask Channel* function was then used to only examine WFA, GAD65/67 and vGLUT1 within this 1.5 μm-wide perimeter surrounding the PV neuron surface.

The presence of WFA labeling in close proximity to the PV neuron surface was assessed for each PV neuron. A PV neuron was considered to have a PNN if there was any WFA labeling around any part of the PV neuron surface as seen by the observer. GAD65/67 and vGLUT1-labeled puncta were then assessed separately using the *Spots* segmentation tool. Within the *Spots* tool, the *Different Spot Sizes (Region Growing)* option was selected and initial settings included an estimated X-Y diameter of 0.5 μm and an estimated Z plane diameter of 0.4 μm. Spots generated by Imaris from these initial settings were then thresholded using the *Classify Spots*, *Quality Filter* histogram to ensure that all labeled puncta were included and background labeling was filtered out. The spots were then thresholded using the *Spot Region, Region Threshold* histogram to ensure that the sizes of the Imaris-generated spots were good approximations of the size of the labeled puncta seen visually by the human observer. Using the *Find Spots Close to Surface Imaris XTension*, we then isolated those spots that were within 0.5 μm of the PV neuron surface. All segmented spots close to the PV neuron surface had to have a Z diameter of at least 0.4 μm to be considered puncta [62, 63].

### Synaptic Electrophysiology

To determine whether there was a change in glutamatergic transmission associated with the nadir (ZT6) and peak (ZT18) periods of PV and PNN staining intensity, a separate cohort of 15 rats was used to measure whole-cell electrophysiology on coronal slices (300 μm collected at 3.2-3.7 mm from bregma) through the prelimbic mPFC. Only one cell per rat was used for each experiment so that reported N-sizes represent the number of animals. Recording conditions and solutions for whole-cell recordings were as previously described [60, 61]. Rats were briefly anesthetized with isoflurane and intracardially perfused with ice cold recovery solution: (in mM) 93 NMDG, 2.5 KCl, 1.2 NaH_2_PO_4_, 30 NaHCO_3_, 20 HEPES, 25 glucose, 4 sodium ascorbate, 2 thiourea, 3 sodium pyruvate, 10 MgSO_4_(H_2_O)_7_, 0.5 CaCl_2_(H_2_O_2_), and HCl added until pH was 7.3-7.4 with an osmolarity of 300-310 mOsm. Slices were prepared on a vibratome (Leica VT1200S) containing recovering solution and then transferred to a holding chamber containing holding solution: (in mM) 92 NaCl, 2.5 KCl, 1.2 NaH_2_PO_4_, 30 NaHCO_3_, 20 HEPES, 25 glucose, 4 sodium ascorbate, 2 thiourea, 3 sodium pyruvate, 2 MgSO_4_(H_2_O)_7_, 2 CaCl_2_(H_2_O_2_), and 2 M NaOH added until pH reached 7.3-7.4 and osmolarity was 300-310 mOsm. Slices remained in the holding chamber for at least 1 h prior to recording. PNN-surrounded fast-spiking interneurons were identified by incubating each slice in holding solution containing FITC-WFA (1 μg/mL) 5 min prior to recording and using CellSens software (Olympus) to identify cells surrounded by fluorescence. The fast-spiking interneurons in this region of the cortex are highly likely to be PV-containing cells [64]. The recording chamber was continuously perfused at 31.0°C at a rate of 4-7 mL/min with artificial cerebrospinal fluid (aCSF): (in mM) 119 NaCl, 2.5 KCl, 1 NaH_2_PO_4_, 26 NaHCO_3_, 11 dextrose, 1.3 MgSO_4_(H_2_O)_7_, and 2.5 CaCl_2_(H_2_O)_2_. Patching pipettes were pulled from borosilicate capillary tubing (Sutter Instruments, CA USA) and the electrode resistance was typically 4-7 mOhms. All experiments utilized cesium chloride (CsCl) internal solution: (in mM) 117 CsCl, 2.8 NaCl, 5 MgCl_2_, 20 HEPES, 2 Mg^2+^ATP, 0.3 Na^2+^GTP, 0.6 EGTA, 0.1 spermine and sucrose to bring osmolarity to 275-280 mOsm and pH to ~7.25. For AMPA:NMDA ratios, neurons were held at +45 mV in 100 μM picrotoxin and 50 μM D-(-)-2-amino-5-phosphonopentanoic acid (d-APV) was added once a stable baseline was acquired. Peak AMPAR excitatory postsynaptic current (EPSC) amplitudes were measured at 20-25 min in d-APV, and this EPSC was subtracted from 5 min averages of baseline EPSCs to obtain the peak NMDAR EPSC [65]. NMDA kinetics were calculated by measuring the time from peak to half peak [66]. All drugs and reagents were obtained from Sigma-Aldrich (St Louis, MO).

### Statistical analysis

For PV, WFA, and 8-oxo-dG staining intensities, distributions of normalized intensities were compared within cell marker between experimental groups using the Kruskal-Wallis test to assess changes among the four ZT times, with all values normalized to ZT0. A Dunn’s multiple comparisons test was used in the case of a significant effect. For comparison among ZTs and each cellular marker (Figure 2), a two-way ANOVA was done and was followed by a post-hoc Bonferroni test in the case of a significant interaction. A Komogorov-Smirnov test was used for comparison of distribution of labeling intensity between two groups. To determine the number of cells expressing each combination of markers, including OTX2, the number of cells was averaged per rat and the results were subjected to a one-way ANOVA followed by a Dunnett’s multiple comparisons test in the case of 4 groups. In cases where comparisons were made between two groups, an unpaired t-test was performed. For puncta analysis, an unpaired t-test was performed or in the case of non-normal distribution, a Mann-Whitney test was performed. TIBCO Software Statistica 13.2 (2016) and GraphPad Prism 8 were used for all intensity, cell number, and puncta analyses. For electrophysiology experiments, unpaired t-tests were used to analyze AMPA:NMDA ratio and NMDA decay rates using Prism 6 (GraphPad Software). Differences were considered significant if p < 0.05.

## RESULTS

### Diurnal fluctuation of PV and PNN intensity

Both PV and WFA intensity fluctuated in a diurnal manner (Fig. 1). Fig. 1A shows an image of single- and double-labeled WFA and PV cells. Fig. 1B and C show cell intensity measures for total WFA+ cells (single-labeled WFA+ cells, including both PV+ and non-PV cells surrounded by WFA) and the double-labeled WFA+/PV+ subset of WFA+ cells. We first tested whether there was a difference between intensities in the light phase (ZT0, ZT6) *vs*. the dark phase (ZT12, ZT18). For WFA+ intensity, there was an effect of the light phase (ZT0, ZT6) *vs*. dark phase (ZT12, ZT18), with increased intensity in the dark phase (p < 0.0001). Fig. 1B shows an effect of ZT (p < 0.0001), with staining intensity of total WFA+ cells increased relative to ZT0: there was a 27% increase at ZT12 (p < 0.0001) and a 40% increase at ZT18 (< 0.0001). Fig. 1C demonstrates a similar pattern for /WFA+/PV+ cells, where there was an effect of the light phase (ZT0, ZT6) *vs*. dark phase (ZT12, ZT18), with increased labeling intensity in the dark phase (p < 0.0001). Fig. 1C also shows that the increases in WFA intensity across the diurnal cycle in WFA+/PV+ double-labeled cells demonstrated an effect of ZT (p < 0.0001) that was similar to that of total WFA+ cells (Fig. 1B), with a 23% increase at ZT12 (p < 0.0001) and a 44% increase at ZT18 (p < 0.0001).

**Figure 1.**
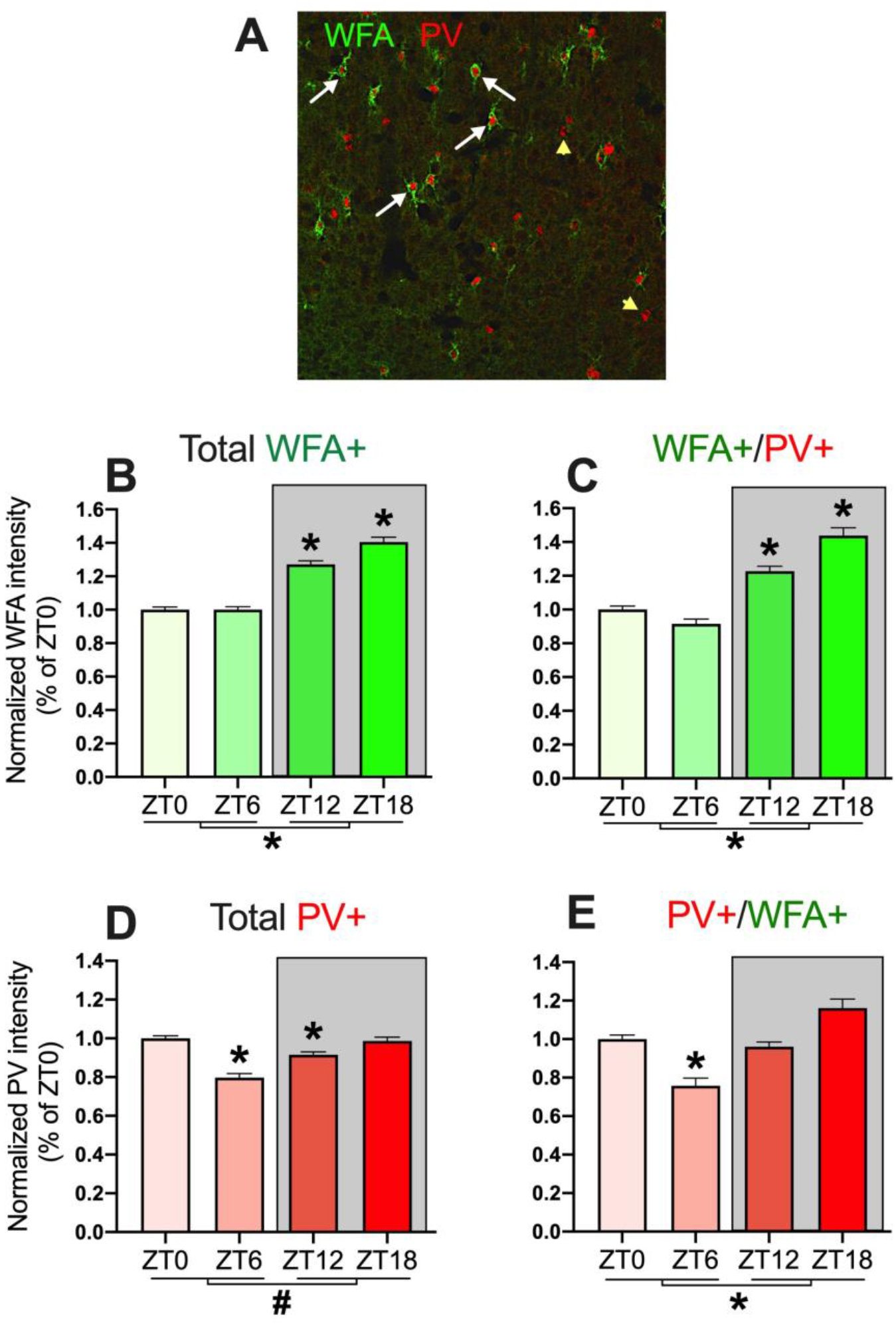
PNN and PV intensity vary across the diurnal cycle. (A) Image (428.81 um^2^) showing WFA+ and PV+ immunolabeling. White arrows are double-labeled cells, yellow arrowheads are single-labeled PV cells (20X). (B) Total WFA intensity in all WFA+ cells was elevated at ZT12 and ZT18 compared with ZT0, and there was a light/dark difference. (C) Similarly, WFA intensity around PV+ cells was elevated at ZT12 and ZT18 compared with ZT0, and there was a light/dark difference. (D) Total PV intensity was decreased in neurons at ZT6 and ZT12 compared with ZT0, and there was a trend for a light/dark difference. (E) PV cell intensity in PV+ cells surrounded by WFA+ was decreased at ZT6 compared with ZT0, and there was a light/dark difference. Data are mean ± SEM; N-size: ZT0, N = 10; ZT6, N = 8; ZT12, N = 7; ZT18, N = 8. *p < 0.05 for individual ZTs compared with ZT0 (individual bars) or for light *vs*. dark comparison (at base of graph); #p < 0.10.

Fig. 1D and E show the patterns of PV labeling intensity in total PV+ cells and those PV+ cells surrounded by WFA (PV+/WFA+). Similar to the WFA studies, we tested whether there was a light *vs*. dark phase difference in PV labeling intensity. Total PV+ cells showed a non-significant but strong trend of the light phase (ZT0, ZT6) *vs*. dark phase (ZT12, ZT18) (p < 0.0624). The labeling intensity of PV+/WFA+ cells showed lower intensity during the light *vs*. dark phase (p < 0.0001). There was an effect of ZT on PV+/WFA+ labeling intensity (p < 0.0001), with a 24% decrease in intensity at ZT6 relative to ZT0 (p < 0.0001).

The number of total WFA+ cells, total PV+ cells, and subsets of cells that were labeled with one or both markers, as well as the percent of co-labeled cells, are presented in Table 1. There was no difference in the number of total WFA+ cells or WFA+/PV-cells across ZT when either comparing the light phase (ZT0, ZT6) with the dark phase (ZT12, ZT18) or when comparing across the four ZTs. The number of total PV+ cells across ZT showed a trend (p = 0.0772). Similarly, the number of WFA+/PV+ cells showed a strong trend (p = 0.0517), with a decrease of about 40% at ZT6 from ZT0 (p = 0.0279). We also examined the percent of WFA+ cells containing PV and *vice versa* and found no overall light *vs*. dark phase effects. For the percent of WFA+ cells that contained PV, there was an effect of ZT (p = 0.022) and a range from 35-58%, with a significant decrease at ZT6 relative to ZT0 (p = 0.011). The percent of PV+ cells surrounded by WFA was consistent across ZTs, ranging from 45-47%. Overall, evaluation of PNNs and PV cells indicate that the intensities of both PNNs and PV cells increased in the dark phase.

**Table 1:**
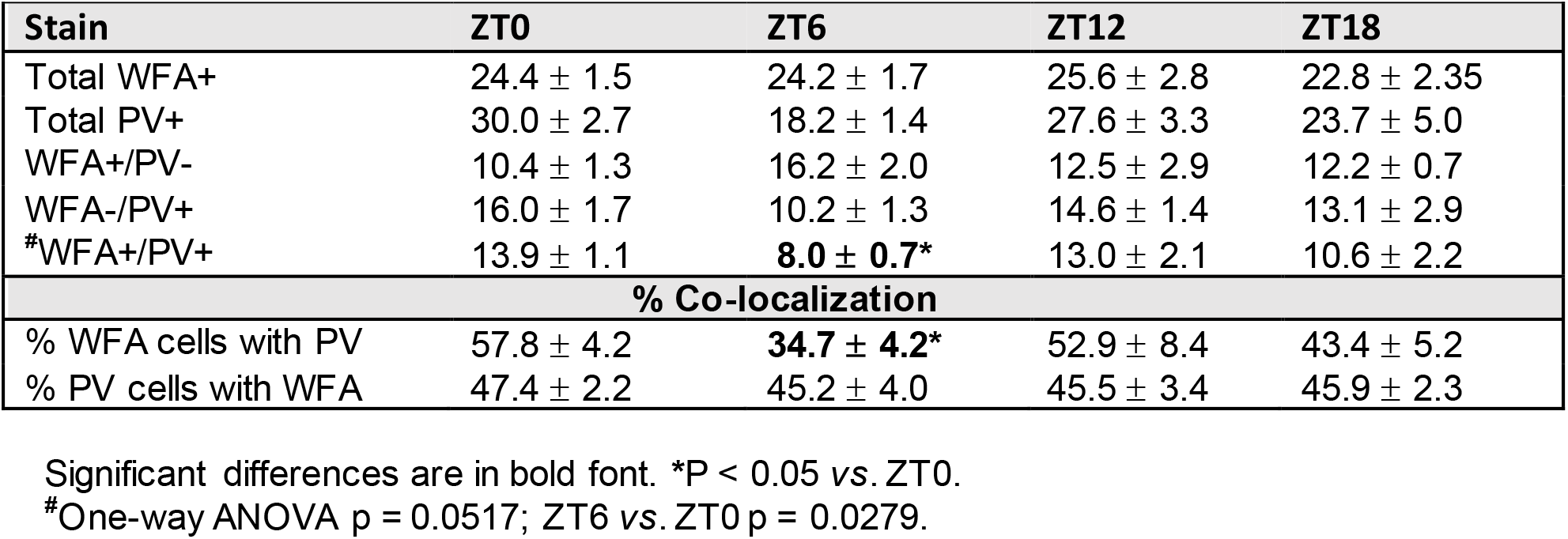
Number and percent of WFA/PV cells.

### Diurnal fluctuation of 8-oxo-dG intensity

We assessed the number of 8-oxo-dG+ cells in a subset of rats shown in Fig. 1. Fig. 2A is shows a region of mPFC containing 8-oxo-dG+, WFA+, and PV+ single, double, and triple-labeled cells. The intensity of total 8-oxo-dG+ cells is shown in Fig. 2B, and we tested whether there was a light *vs.* dark phase difference. There was an effect of the light phase (ZT0, ZT6) *vs*. dark phase (ZT12, ZT18), with decreased intensity in the dark phase (p < 0.0001). There was an effect of ZT on 8-oxo-dG intensity (p < 0.0001), with a decrease at all ZTs relative to ZT0 (p < 0.0001 for all ZTs), with a maximal 47% decrease at ZT12 *vs*. ZT0. The significant decrease in intensity of 8-oxo-dG was maintained at ZT18, with a 27% decrease *vs*. ZT0. Fig. 2C shows the intensity of 8-oxo-dG in 8-oxo-dG+/WFA+/PV+ triple-labeled cells, with an effect of light *vs*. dark phase (p < 0.0001) and an effect of ZT (p < 0.0001). These triple-labeled cells showed a 20-25% decrease in intensity of 8-oxo-dG at all three ZTs relative to ZT0 (p < 0.0001 for all ZTs). Comparison of total 8-oxo-dG+ cells with 8-oxo-dG+/WFA+/PV+ triple-labeled cells showed a main effect of cellular subtype (single-*vs*. triple-label) across ZTs (F_1,22,121_ = 133.5, p < 0.0001), a main effect of ZT (_F2,22,121_ = 38.6, p < 0.0001), and a cell subtype × ZT interaction (_F2,22,121_ = 10.23, P < 0.0001), with an increase in 8-oxo-dG intensity in triple *vs.* total 8-oxo-dG+ cells at all ZTs (p < 0.0001 for both; not shown).

**Figure 2.**
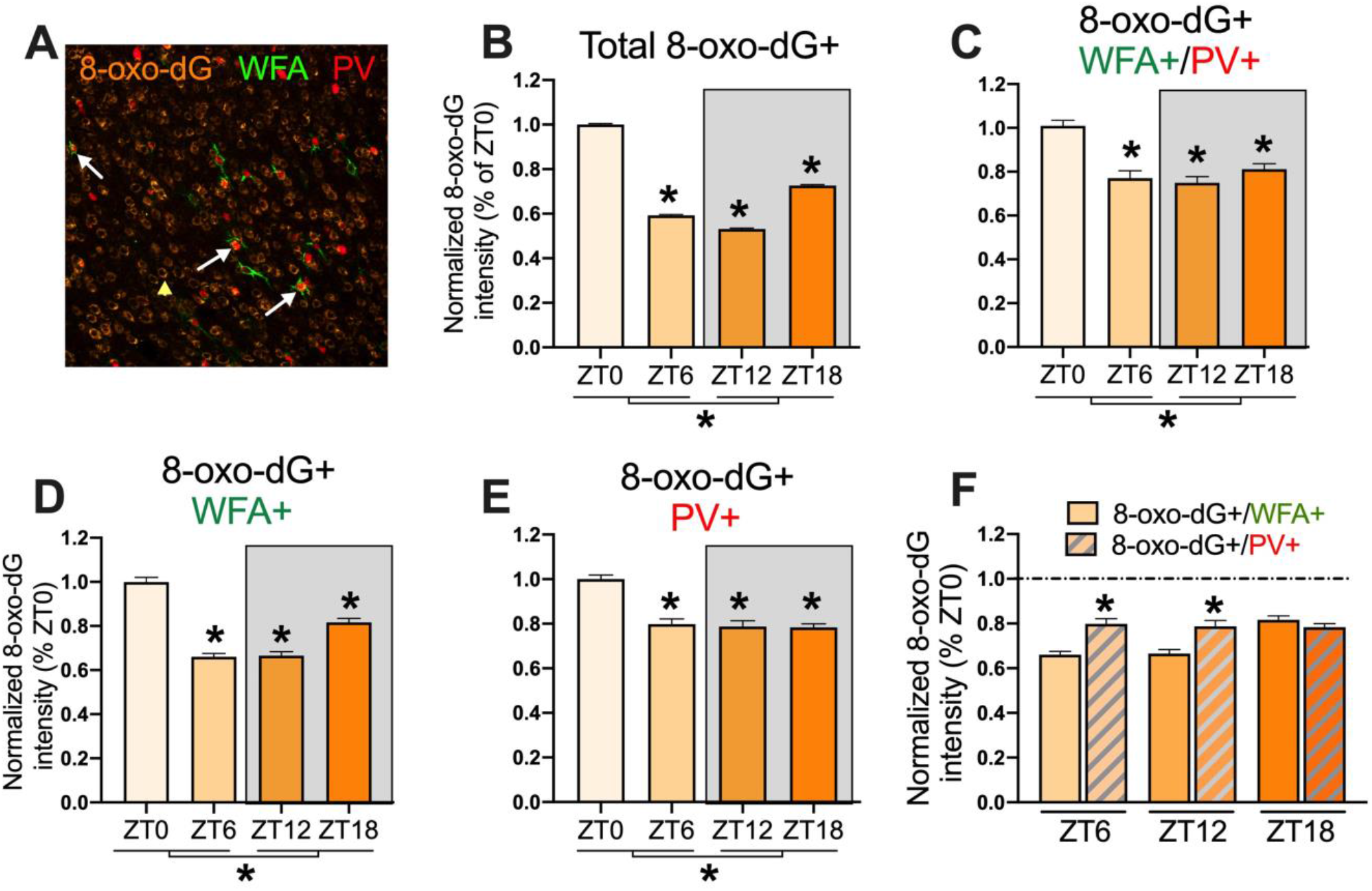
The oxidative stress marker 8-oxo-dG varies in PV cells with or without PNNs across the diurnal cycle. **(A)** Image (428.81 um^2^) showing 8-oxo-dG+, WFA+, and PV+ immunolabeling. White arrows are triple-labeled cells; yellow arrow is single-labeled 8-oxo-dG cell (20X). (B) Total 8-oxo-dG was decreased at all ZTs compared with ZT0, and there was a light/dark difference. (C) Triple-labeled 8-oxo-dG+/WFA+/PV+ cells demonstrated a decrease at all ZTs compared with ZT0, and there was a light/dark difference. (D) Double-labeled 8-oxo-dG+/WFA+ cells demonstrated a decrease at all ZTs compared with ZT0, and there was a light/dark difference. (E) Double-labeled 8-oxo-dG+/PV+ cells demonstrated a decrease at all ZTs compared with ZT0, and there was a light/dark difference. (F) 8-oxo-dG+/PV+ cells showed higher oxidative stress levels than 8-oxo-dG+/WFA+ cells at ZT6 and ZT12. Each group of double-labeled cells was normalized to its own ZT0 value; dotted line is normalized ZT0 value. Data are mean ± SEM; N-size: ZT0, N = 4; ZT6, N = 4; ZT12, N = 3; ZT18, N = 4. For B-E, *p < 0.05 compared with ZT0 for individual ZTs compared with ZT0 (individual bars) or for light *vs*. dark comparison (at base of graph); for (F), *p < 0.05 compared with 8-oxo-dG/WFA+ cells within the same ZT.

This increase in intensity of 8-oxo-dG in triple-labeled cells may be due to either the presence of PNNs, the presence of PV, or both, so we separately analyzed the intensity of 8-oxo-dG in 8-oxo-dG+/WFA+ cells and 8-oxo-dG+/PV+ cells across ZTs. The intensity of 8-oxo-dG in 8-oxo-dG+/WFA+ cells shown in Fig. 2D indicates an effect of light *vs.* dark phase (p < 0.0001) as well as an effect of ZT (p < 0.0001). As with single-and triple-labeled cells, the intensity of 8-oxo-dG in cells with WFA+ was reduced at all ZTs relative to ZT0 (p < 0.0001 for all ZTs). The intensity of 8-oxo-dG in 8-oxo-DG+/PV+ cells shown in Fig. 2E indicates an effect of light *vs*. dark phase (p < 0.0001) as well as an effect of ZT (p < 0.0001). The intensity of 8-oxo-dG in PV cells was reduced at all ZTs relative to ZT0 (p < 0.0001 for all ZTs). We then compared the intensity of 8-oxo-dG+/WFA+ *vs*. 8-oxo-dG+/PV+ cells (Fig. 2F). There was a main effect of cell subtype (F_1,3299_ = 23.9, p < 0.0001), a main effect of ZT (F_2,3299_ = 12.4, p < 0.0001), and a cell subtype × ZT interaction (F_1,3299_ = 15.7, p < 0.0001), with greater increases in 8-oxo-dG intensity in PV+ containing cells compared with WFA+ containing cells. Therefore, a major determinant of 8-oxo-dG buildup was the PV cell phenotype rather than the presence of PNNs.

We analyzed whether the magnitude of 8-oxo-dG intensity was associated with WFA or PV intensity. We conducted a median-split analysis of 8-oxo-dG intensity for all ZTs combined, dividing 8-oxo-dG into low or high intensity. We then examined whether WFA intensity (expressed as percent of its own ZT0 as before) was different in the low-vs. high-intensity 8-oxo-dG-containing cells. WFA intensity in low-intensity cells was 97.2 ± 2.5% and in high-intensity cells was 137.0 ± 4.5% (p < 0.0001). PV intensity in low-intensity was 64.7 ± 1.9% and in high-intensity cells was 95 ± 2.4% (p < 0.0001). Therefore, in 8-oxo-dG+/PV+/WFA+ cells, both WFA intensity and PV intensity were higher in cells with high 8-oxo-dG intensity than in cells with low 8-oxo-dG intensity.

We also examined the number and percent of 8-oxo-dG+ single, double, and triple-labeled cells (Table 2). There was an effect of ZT only for 8-oxo-dG+/PV+ double-labeled cells (p = 0.007), with a decrease of approximately 50% at ZT6 (p = 0.008) and a decrease of approximately 40% at ZT12 (p = 0.037). There was also an effect of ZT for 8-oxo-dG+/WFA+/PV+ triple-labeled cells (p = 0.021), with a decrease of approximately 60% at ZT6 relative to ZT0 (p = 0.015). For the percent of 8-oxo-dG double-and triple-labeled cells, only 5-9% of these cells contained either WFA or PV, and about 2-4% contained both WFA and PV. The vast majority of WFA+ cells contained 8-oxo-dG (86-92%), and, as with WFA+ single-labeled cells, there was an effect of light *vs.* dark phase for the percent of WFA+ cells positive for 8-oxo-dG (p = 0.010). Similarly, most WFA+/PV+ neurons also contained 8-oxo-dG (93-96%). In contrast, only 56-59% of PV+ cells contained 8-oxo-dG, suggesting that there was a substantial (40%) population of PV neurons that may be much less metabolically active compared with those surrounded by PNNs or in which 8-oxo-dG was too low to be detected in our system. Overall, these results indicate that oxidative stress levels, as indicated by 8-oxo-dG levels, is highest in PV+ cells, and the diurnal decrease in 8-oxo-dG levels at ZT6 and ZT12 is larger in non-PV+ cells compared with PV+ cells.

**Table 2:** Number and percent of 8-oxo-dG cells.

| Stain | ZT0 | ZT6 | ZT12 | ZT18 |
| --- | --- | --- | --- | --- |
| Total 8-oxo-dG+ | 333.9 ± 14.7 | 295.5 ± 50.8 | 330.7 ± 33.0 | 339.1 ± 30.5 |
| 8-oxo-dG+/WFA+ | 24.5 ± 1.4 | 23.0 ± 1.4 | 28.6 ± 2.5 | 24.6 ± 3.0 |
| 8-oxo-dG+/WFA- | 310.7 ± 14.6 | 272.5 ± 48.7 | 298.4 ± 35.5 | 314.5 ± 28.0 |
| 8-oxo-dG+/PV+ | 23.8 ± 2.2 | **11.8 ± 1.5\*** | **13.6 ± 3.4\*** | 22.4 ± 2.1 |
| 8-oxo-dG+/PV- | 311.3 ± 13.3 | 283.7 ± 51.6 | 313.3 ± 36.4 | 316.7 ± 28.8 |
| 8-oxo-dG+/WFA+/PV+ | 15.4 ± 0.7 | **6.3 ± 0.5\*** | 10.3 ± 3.7 | 14.5 ± 2.2 |
| **% Co-localization** |  |  |  |  |
| **% of 8-oxo-dG+ cells** |  |  |  |  |
| with WFA+ | 7.3 ± 0.5 | 8.1 ± 0.7 | 9.1 ± 1.5 | 7.2 ± 0.4 |
| with PV+ | 7.0 ± 0.5 | 4.5 ± 1.1 | 4.6 ± 1.4 | 6.6 ± 0.4 |
| with WFA+/PV+ | 4.5 ± 0.1 | 2.3 ± 0.4 | 3.5 ± 1.4 | 4.2 ± 0.5 |
| **% of WFA+ cells** |  |  |  |  |
| with 8-oxo-dG+ | **87.0 ± 1.3<sup>1</sup>** | **85.5 ± 1.9<sup>1</sup>** | **91.6 ± 0.4<sup>2</sup>** | **90.0 ± 1.6<sup>2</sup>** |
| **% of PV+ cells** |  |  |  |  |
| with 8-oxo-dG+ | 59.3 ± 4.9 | 56.9 ± 1.1 | 55.5 ± 4.3 | 58.2 ± 4.1 |
| **% of WFA+/PV+ cells** |  |  |  |  |
| with 8-oxo-dG | 94.7 ± 1.3 | 93.1 ± 1.6 | 93.7 ± 2.5 | 96.2 ± 0.1 |

Significant differences are in bold font. Numbers <sup>1,2</sup> denote difference between light (ZT0 + ZT6) and dark (ZT12 + ZT18). \*P < 0.05 from ZT0.

### Diurnal pattern of excitatory puncta in WFA+/PV+ cells at ZT6 vs. ZT18

We observed the largest differences in intensity of WFA+/PV+ staining between ZT6 and ZT18 and therefore chose these two time points to further test whether there were diurnal differences in inhibitory and excitatory inputs to these cells. Fig. 3A and 3B show a representative PV neuron surrounded by a WFA-labeled PNN and apposed by GAD65/67-labeled inhibitory and vGLUT1-labeled excitatory puncta. While the number of GAD65/67 puncta was not different between ZT6 and ZT18 (Fig. 3C), the number of vGLUT1 puncta was increased at ZT18 compared with ZT6 (p = 0.0075; Fig. 3D). There was a strong trend for a decreased ratio of GAD65/67:vGLUT1 at ZT18 compared with ZT6 (p = 0.0529; Fig. 3E) and a small but significant increase in the volume of the rendered surface through the middle of the analyzed WFA+/PV+ cells (p = 0.0236; Fig. 3F).

**Figure 3.**
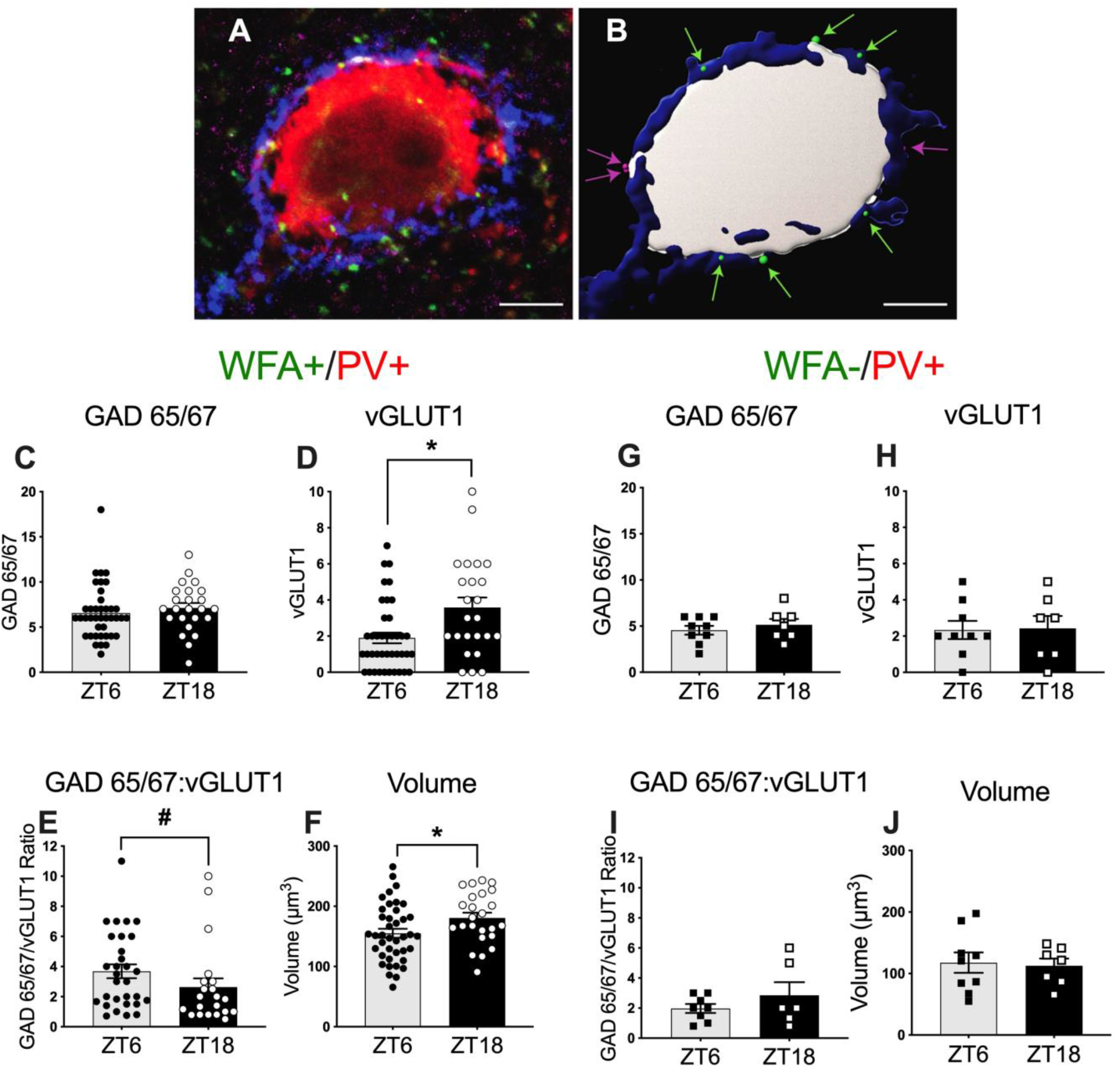
The number of glutamatergic puncta apposing PV neurons with PNNs increases at ZT18, and PV cell volume increases at ZT18. (A) An immunolabeled PV neuron (red) is surrounded by a WFA-labeled PNN (blue) in a representative confocal micrograph from the mPFC. The PV neuron is receiving appositions from both vGLUT1-labeled glutamatergic puncta (magenta) and GAD65/67-labeled GABAergic puncta (green). (B) The Imaris *Surfaces* segmentation tool was used to render the PV neuron (gray) and WFA-labeled PNN (blue). The Imaris *Spots* segmentation tool was used to segment GAD65/67 (green arrows) and vGLUT1-labeled (magenta arrows) puncta that met size and location criteria. Scale bar = 5 μm. **In WFA+/PV+ cells:** (C) Number of inhibitory puncta (GAD65/67) were similar at ZT6 and ZT18. (D) Number of excitatory puncta (vGLUT1) apposing PV neurons was higher at ZT18 compared with ZT6. (E) The ratio of GAD65/67:vGLUT1 puncta trended toward a decrease at ZT18 compared with ZT6. (F) The volumes of the three-dimensional surfaces rendered with Imaris software through the middle of the PV neurons were larger at ZT18 compared with those at ZT6. **In WFA-/PV+ cells:** (G) Number of inhibitory puncta (GAD65/67) was similar in the ZT6 and ZT18 groups. (H) Number of excitatory puncta (vGLUT1) was similar at ZT6 and ZT18. (I) The ratio of GAD65/67:vGLUT1 puncta was similar at ZT6 and ZT18. (J) The volumes of the three-dimensional surfaces rendered with Imaris software were similar at ZT6 and ZT18. Data are mean ± SEM; N = 4/group. *p < 0.05 compared with ZT6; #p < 0.10 compared with ZT6.

We also measured the number of GAD65/67 and vGLUT1 puncta apposing PV cells devoid of PNNs (WFA-/PV+ cells) to determine whether there were diurnal differences similar to those found in PNN-surrounded PV cells. Fig. 3G-J show that there were no differences in inhibitory or excitatory puncta or in the volume of WFA-/PV+ cells. The volume of WFA+/PV+ cells was significantly larger than WFA-/PV+ cells (WFA+ effect F_1,76_ = 16.79; p < 0.0001), as previously reported for humans [67], and the volume of PV cells was positively correlated with PNN intensity (p = 0.0237; data not shown). Overall, these findings suggest that PV cell volume and diurnal changes in excitatory puncta are dependent on the presence of PNNs.

### Diurnal pattern of AMPA:NMDA ratio in WFA+/PV+ cells at ZT6 vs. ZT18

To determine whether there may also be post-synaptic changes in excitatory transmission at ZT6 and ZT18 we measured the AMPA:NMDA ratio in WFA+ fast-spiking interneurons at these two time points. The fast-spiking interneurons in this region of the cortex are highly likely to be PV-containing cells [64]. Fig. 4A shows a representative trace of the amplitudes from AMPA- and NMDA-mediated currents. The individual amplitudes for AMPA EPSCs were greater at ZT18 compared with ZT6, and the AMPA:NMDA ratio of WFA+/PV+ cells shown in Fig. 4B increased at ZT18 compared with ZT6 (ZT6 N = 7: 0.55 ± 0.05; ZT18 N = 8: 0.91 ± 0.13, p = 0.0347). This increase in AMPA:NMDA ratio could be due to increased calcium-permeable (CP)-AMPAR utilization, decreased NMDA-mediated transmission, or a combination of both. Additionally, we observed an apparent difference in the slopes of the isolated NMDAR current (Fig. 4A, green traces). Further evaluation of NMDAR decay kinetics shown in Fig. 4C revealed a trending increase in the time constant at ZT18 compared with ZT6, a common indicator of a shift in subunit composition from GluN2A (highly expressed on PV+ fast-spiking cells) to GluN2B (ZT6 N = 7: 41.63 ms ± 6.27; ZT18 N = 8: 54.74 ms ± 4.12, p = 0.0965).

**Figure 4.**
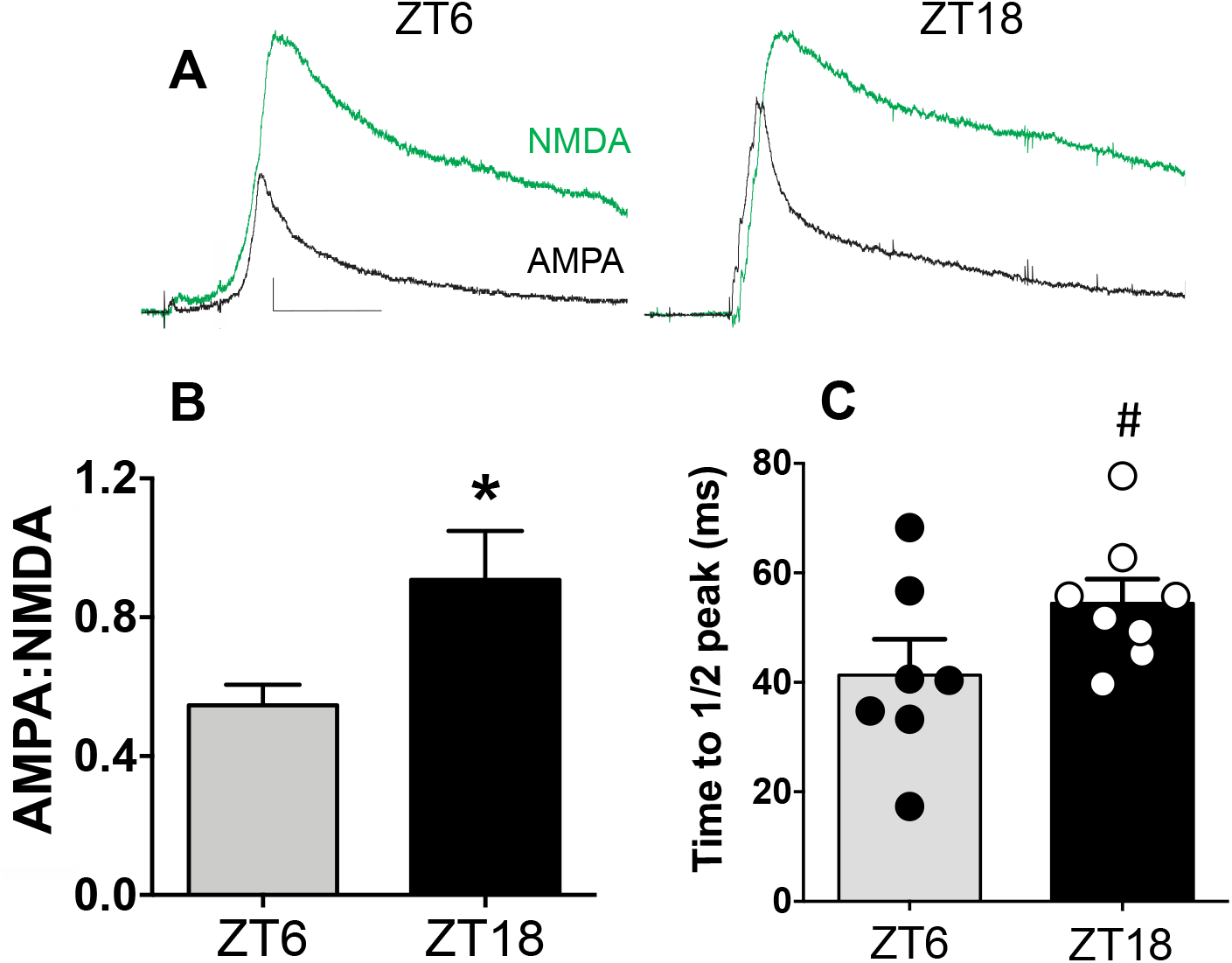
The AMPA:NMDA ratio increases at ZT18. (A) Representative traces of the AMPA (black) and NMDA (green) components at ZT6 and ZT18. Scale bar represents 10 pA, 10ms. (B) AMPA:NMDA ratio of mEPSCs evoked from PNN-surrounded fast-spiking cells in the prelimbic mPFC increases during the dark phase at ZT18. (C) Average time constants of decay of the NMDA component of mEPSCs evoked from PNN surrounded PV FSIs in the prelimbic mPFC. Data are mean ± SEM; Number of rats: ZT6 N = 7; ZT18 N = 8. *p < 0.05; #p < 0.10 compared with ZT6.

### Number of OTX2+ cells at ZT6 vs. ZT18

Because OTX2 is necessary to maintain PNNs [52–55], we also determined whether there was a diurnal difference in the number of WFA+/PV+ cells double- or triple-labeled with OTX2 between ZT6 and ZT18. Fig. 5A shows a region of mPFC containing single-, double-, and triple-labeled OTX2+ cells. Fig. 5B shows that there were no differences across timepoints in the total number of cells labeled with OTX2+ or in number of OTX2+/WFA+ cells, whereas there was a strong trend for an increased number of double-labeled OTX2+/PV+ cells (p = 0.0609) and triple-labeled OTX2+/WFA+/PV+ cells (p = 0.0770). Interestingly, the number of PV cells that did not co-label with OTX2+ was not different between ZT6 and ZT18 (p = 0.1936), suggesting that PV+/OTX2-cells may constitute a different PV phenotype that is not regulated diurnally.

**Figure 5.**
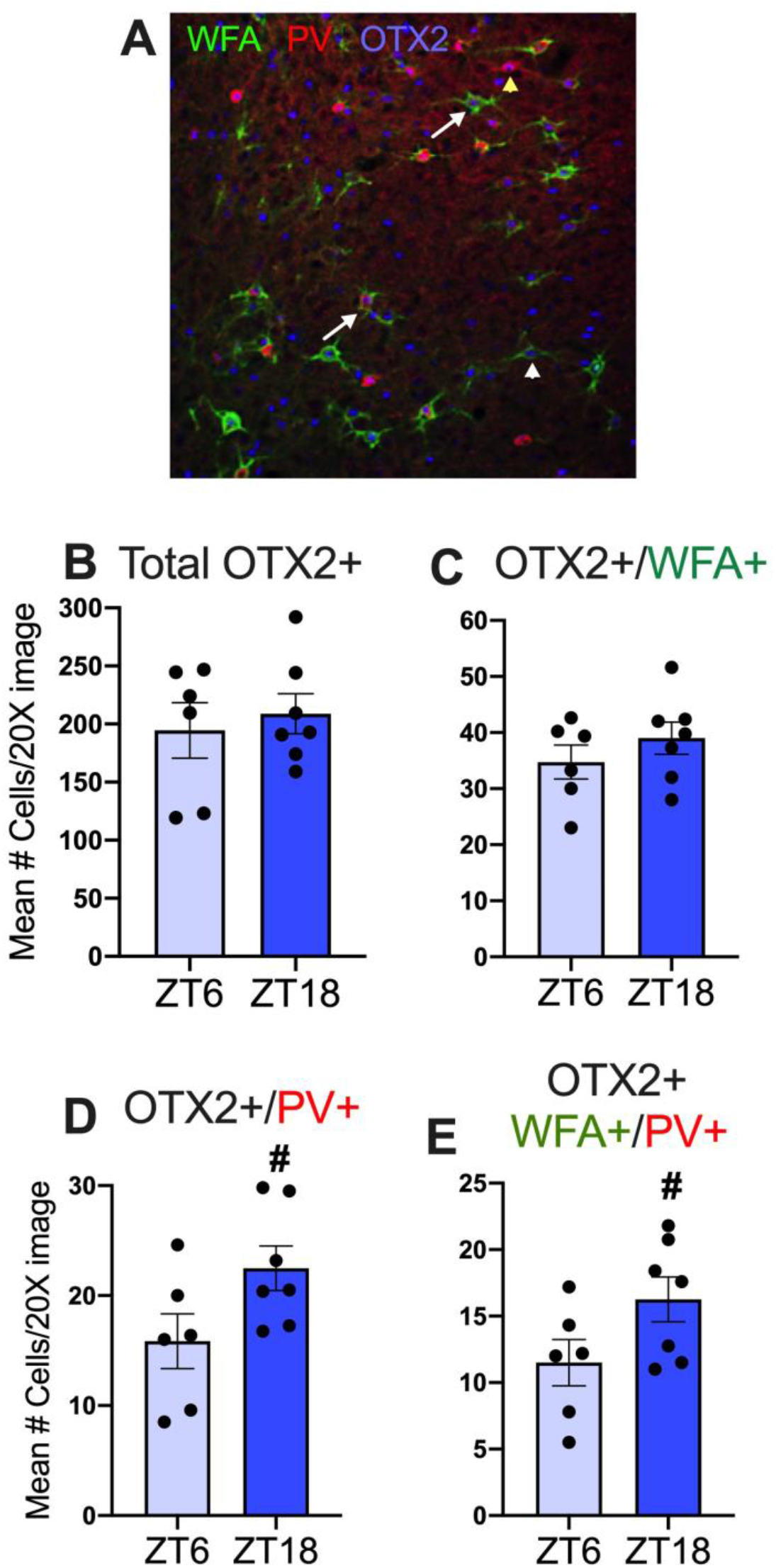
OTX2 staining in PV+ cells and WFA+/PV+ cells trend toward increase at ZT18. (A) Image (428.81 um^2^) showing OTX2+, WFA+, and PV+ immunolabeling. White arrows are triple-labeled cells, yellow arrowhead is double-labeled OTX2+/PV+ cell, and white arrowhead is double-labeled OTX2/WFA+ cell (20X). (B) Total number of OTX2+ cells was similar at ZT6 and ZT18. (C) Number of OTX2+/WFA+ cells was similar at ZT6 and ZT18. (D) Number of OTX2+/PV+ cells showed a strong trend toward increase at ZT18 compared with ZT6. (E) Number of OTX2+/WFA+/PV+ cells showed a trend toward increase at ZT18 compared with ZT6. Data are mean ± SEM; N = 6-7/group. #p < 0.10 compared with ZT6.

The number of OTX2+ single-, double-, and triple-labeled cells and the percent of co-labeled cells was also determined (Table 3). There were no other changes in the number of these cells between ZT6 and ZT18 beyond those shown in Fig. 5. Approximately 20% of OTX2+ cells were co-labeled with WFA, whereas only about 10% of OTX2+ cells were co-labeled with PV, and about 7% were co-labeled with both WFA and PV. The majority of cells staining for WFA or PV or both WFA and PV were co-labeled with OTX2 (ranging from 67-94%). Overall, these findings indicate that OTX2 is not limited to a single PV-containing phenotype, and the number of OTX2 cells in PV neurons trends toward an increase at ZT18 *vs.* ZT6.

**Table 3:**
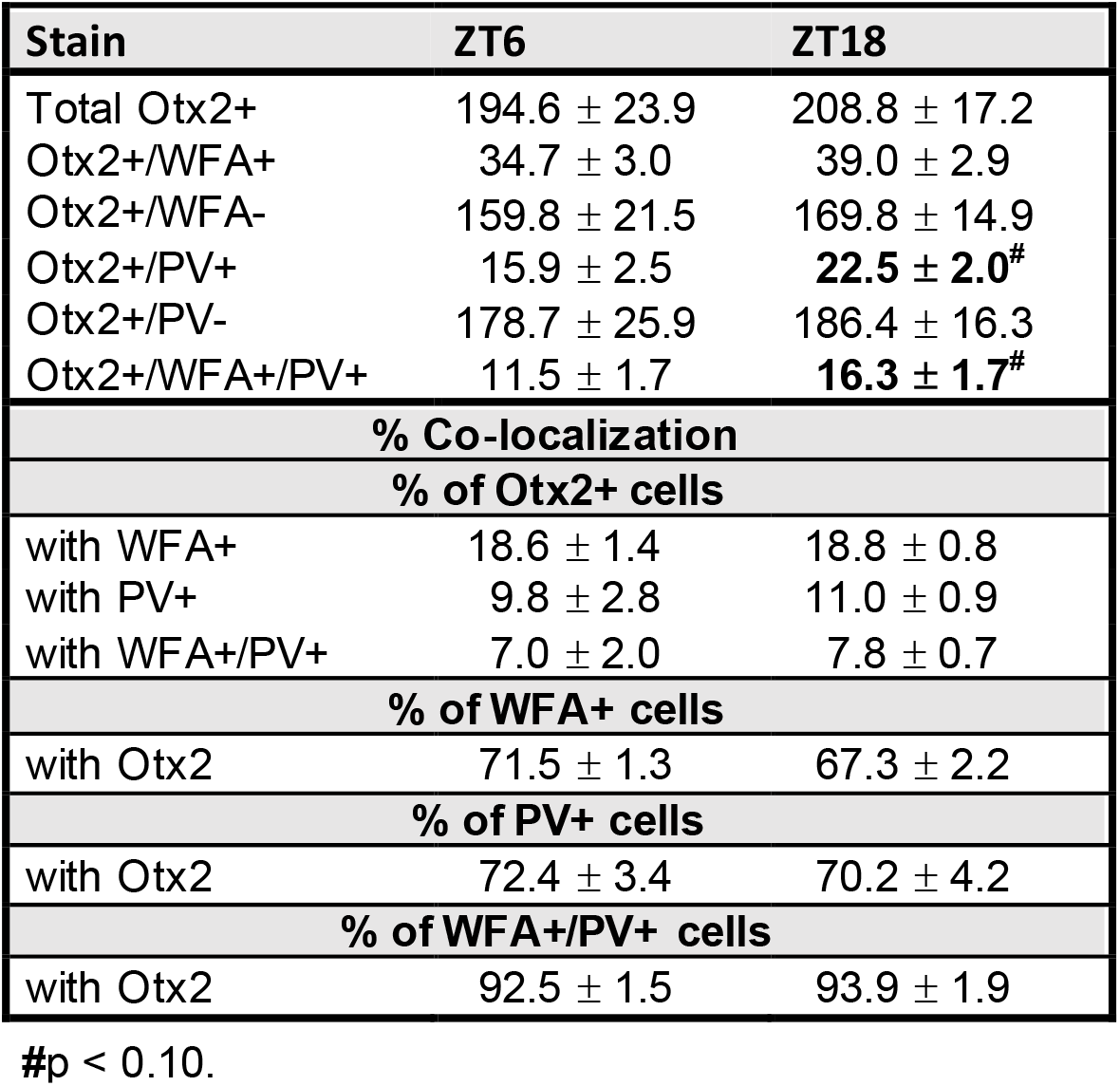
Number and percent of Otx2 cells.

## DISCUSSION

Here we report several novel findings regarding diurnal variations in the biochemical features and physiology of PV cells and PNNs: 1) the intensities of biochemical markers for PV cells and PNNs in the mPFC increased during the dark, active period; 2) the oxidative stress marker, 8-oxo-dG, decreased across all time points relative to ZT0, with decreases most pronounced in non-PV-containing cells; 3) the increase in PV and PNN intensity at ZT18 were accompanied by an increase in excitatory (vGLUT1) puncta in PV cells surrounded by PNNs and, in parallel with this finding, 4) the AMPA:NMDA ratio was increased in PV/PNN cells; and 5) the number of cells co-labeled for PV/OTX2 and PV/WFA/OTX2 showed strong trends toward increases at ZT18 relative to ZT6.

### PNN intensity is increased in the dark phase

The intensity of PNNs was higher in the dark phase (ZT12, ZT18) compared with the light phase (ZT0, ZT6). Diurnal fluctuation in the number of WFA-labeled PNNs has recently been reported for mice and humans [49]. The study in mice revealed that the number of PNNs in several brain regions, including in the mPFC, followed a diurnal pattern, with higher numbers of PNNs in the dark, active phase. PNN-surrounded neurons in mPFC also demonstrated a similar diurnal pattern when mice were held under conditions of constant darkness, indicating the presence of a true circadian rhythm [49]. A circadian contribution (*vs*. sleep contribution) to the diurnal rhythm is consistent with the minimal changes in PNN intensity we previously observed at ZT6 after 6 hr sleep disruption [48]. In contrast to the Pantazopoulos study in mice [49], we found no diurnal rhythm in the *number* of WFA-labeled PNNs but instead a diurnal rhythm in the *intensity* of PNNs, suggesting that perhaps very dim PNNs were not visible in the mouse study or that there are species differences in PNN expression. Importantly, the intensity of PNNs influences the extent of plasticity of their underlying neurons, based on studies in knockout mice missing a critical proteoglycan for PNN formation (link protein 1); these mice have much less intense PNNs yet exhibit delayed critical period plasticity in the visual cortex [2]. In studies in which PNNs are removed by Ch-ABC, decreased intensity of PNNs in the light phase would be expected to impart an increase in membrane capacitance [68], a decrease in firing rate [4, 68, 69], a lower resting membrane potential [70] and changes in ion mobility around PV cells, including Ca^2+^ [71, 72]. Thus, an increase in PNN intensity during the active phase may allow for underlying neurons to maintain high firing rates when sleep is less likely to occur, since removal of PNNs decreases the firing rate of PV neurons. In turn, a decrease in PNN intensity in the inactive phase may promote high plasticity during sleep in the light phase, leading to stabilization or removal of synapses needed for memory consolidation [73–76].

The particular component(s) of PNNs that are diurnally regulated is unknown. The daily decrease in PNNs appears to be regulated at least in part by cathepsin-S in the mPFC [49]. Pantazopoulos *et al*. [49] recently suggested that circadian changes in the number of PNNs may be related to changes in the sulfation pattern on chondroitin sulfate proteoglycans of PNNs [49], since it is difficult to conceive that the entirety of PNNs would fluctuate diurnally. Two key sulfation patterns on chondroitin sulfate chains attached to proteoglycans in PNNs are the 4-sulfation and 6-sulfation pattern, with 4-sulfation chondroitin sulfate chains prevalent in adults [2, 53]. WFA appears to bind to the 4-sulfated chains on aggrecan [26, 77], suggesting that aggrecan or sulfation patterns on aggrecan may be reduced during the light period and restored during the more active dark period. But how these changes would occur on a rapid, daily basis is unknown. The composition of chondroitin sulfate chains is central to regulating plasticity because these chains form binding sites for several molecules on PNNs, including semaphorin 3A [78], neuronal-activity regulated pentraxin 2 (Nptx2), [79], which clusters AMPARs on PV neurons [80], and the transcription factor OTX2 [52, 53, 55]; all of these molecules regulate plasticity.

### OTX2 expression in the light vs. dark phase

We examined whether there was diurnal expression of OTX2, which is synthesized in the choroid plexus [81], binds to PNNs [52], and is internalized within PNN-surrounded cells where it acts as a transcription factor [52, 54]. OTX2 is transferred to PV cells in the visual cortex and is responsible for both opening and closing the critical period of plasticity in part through its binding to PNNs [52, 55]. In the prelimbic mPFC, about 20% of OTX2-containing cells co-expressed PNNs (Table 3). Unlike in the visual cortex [52], OTX2 in the mPFC is not preferentially accumulated by WFA+ or PV+ cells ([82]; this study). On the other hand, the majority of single-labeled WFA+ cells, PV+ cells, and double-labeled WFA+/PV+ cells contained OTX2. While we found no diurnal difference in the *total* number of cells expressing OTX2, we did observe a strong trend toward diurnal rhythmicity in both the number of *PV+ cells* and *WFA+/PV+ cells* containing OTX2 (Figure 5; Table 3). This finding suggests that OTX2 may accumulate specifically in PV+ cells in a diurnal-dependent manner to upregulate PV and PNNs and in turn regulate excitatory:inhibitory balance. Diurnal expression of *OTX2* mRNA has been demonstrated in the rat pineal gland, with maximal expression in the dark phase [83]. OTX2 has been suggested to reciprocally interact with CLOCK in a positive regulatory loop in Xenopus embryos [84], and analysis of OTX2-dependent critical period gene expression identified upregulation of *Per1* in mouse visual cortex [85].

OTX2 regulates several PNN components in the mPFC, possibly by altering turnover of these components [82]. Several downstream targets of OTX2 have been identified, including the GluN2A subunit of the NMDA receptor (see below), and the antioxidant gene oxidation resistance 1 (*Oxr1)*, whose gene product acts as a sensor of oxidative stress and is enriched in GABAergic neurons in the visual cortex [50]. OTX2 also may bind to several potassium channel genes, including the Kv3.1 family, a key ion channel that maintains fast spiking in PV cells [86]. Thus, diurnal expression of OTX2 likely coordinates the expression of several important genes in PV cells to render these neurons responsive to external signals [50] and oxidative stress to support daily fluctuation of firing rates and gamma oscillations needed to provide appropriate excitatory:inhibitory balance during sleep and wakefulness [87].

### PV cell intensity, excitatory puncta, and AMPA:NMDA ratio are increased in the dark phase

We also demonstrated diurnal rhythmicity of PV expression in mPFC cells. PV protein levels have shown a circadian pattern of expression in retinal amacrine cells, with higher expression in the dark, active phase [88]. Expression of PV is regulated by *Clock* gene expression in the visual cortex [89]. Thus, similar CLOCK-dependent rhythms of PV levels may occur in the prelimbic mPFC. Diurnal changes in PV intensity have been shown to be positively correlated with GAD67 intensity [90] and likely reflect activity levels needed to sustain homeostatic balance for excitatory:inhibitory output. Parvalbumin is a Ca^2+^ buffering protein [91], which may lead to multi-faceted consequences of diurnal variation in PV levels. For example, low expression of PV in the absence of prolonged bursts of action potentials may promote short-term plasticity through facilitation [92, 93], while high expression of PV may lower the release probability of GABA by acting as a fast Ca^2+^ buffer [93]. Low PV levels may therefore function to amplify low-input signals, whereas high PV levels may function as a high-pass filter to dampen low-input signals. High PV levels may also act as a brake on excitation-transcription coupling *via* a cAMP response element-binding protein (CREB)-dependent process [94], which in turn regulates several genes [95] that could orchestrate day/night fluctuations in PV cell function.

In concert with increases in PV levels during the dark phase, we found two additional diurnal changes consistent with increases in glutamate transmission at ZT18. First, the AMPA:NMDA ratio was increased in WFA+/PV+ cells at ZT18 relative to ZT6, which may drive a higher cell firing rate. Previous observations in neurons of the cerebral cortex indicated that protracted wakefulness upregulates AMPA receptor-mediated signaling [96], but the neurochemical phenotype of these neurons is unknown. Therefore, the increase in AMPA/NMDA ratio we observed at ZT18 (a time when rats have been awake for several hours after dark onset) may be driven by sustained wakefulness rather than an endogenous circadian oscillator. In PV cells, AMPA receptors have an abundance of GluA1 and GluA4 subunits, making them calcium permeable [97], which contributes to faster kinetic properties, including higher single-channel conductance and faster gating properties compared with GluA2-containing AMPARs [97, 98]. Second, we observed an increase in the number of vGLUT1 synaptic puncta apposing WFA+/PV+ cells at ZT18 relative to ZT6 (Fig. 3). Interestingly, we found no changes in vGLUT1 puncta apposing PV+ cells *devoid* of PNNs (WFA-/PV+ cells), suggesting that the presumed higher firing rate of PV cells with PNNs *vs*. those without PNNs may contribute to maintaining diurnal fluctuation in excitatory:inhibitory balance. It is also possible that NMDARs are altered diurnally. Our recordings demonstrated that the NMDA current was greater than the AMPA current. There is an early switch during postnatal development from GluN2B to GluN2A expression [99], and PV function is maintained by GluN2A-containing NMDA receptors [100]. The trend toward a slower time constant in PV+/WFA+ cells at ZT18 suggests that the NMDA component is more NR2B dominant, as slower time constants indicate a shift from GluN2A to GluN2B composition [66]. Future experiments are needed to determine the extent of diurnal rhythm-induced compositional changes in NMDA receptors. Overall, our findings indicate that glutamate transmission is high during the active phase at ZT18 relative to the inactive phase at ZT6. The reduction in PNN intensity during the inactive phase may lead to lower CP-AMPARs in part *via* reduced accumulation of Nptx2 within PNN-surrounded PV cells [101]. Collectively, these changes may help coordinate not only excitatory:inhibitory balance needed for pyramidal cell output during wakefulness, but also for homeosynaptic scaling during sleep to restore synaptic homeostasis [74]. Importantly, most rodent studies are conducted in the daytime phase, and thus modulation of PV cell electrophysiology and transcription-coupling by factors such as glutamatergic input that regulates pCREB in pyramidal cells but not in PV cells [94] might be unveiled if PV cells were tested during the dark phase.

### Oxidative stress is highest at ZT0 and remains higher in PV vs. non-PV cells

Diurnal changes in oxidative stress are observed in a multitude of studies, demonstrated by fluctuations in the extent of DNA damage, lipid peroxidation, and protein oxidation [102, 103]. These downstream effects appear to be initiated by the impact of accumulated reactive oxygen species (ROS) [102, 104]. The diurnal oscillation we observed in the oxidative stress marker 8-oxo-dG aligns with our previous findings [48] and previous studies [105, 106] that oxidative stress accumulates in rodent brains during wakefulness. The diurnal changes reflect the rhythmicity of mitochondrial network remodeling and the dynamic capacity of oxidative phosphorylation, which peaks in the dark phase [107] during wakefulness relative to sleep (reviewed in [108, 109]). Here we show that 8-oxo-dG intensity in the prelimbic mPFC was highest at ZT0 and lowest at ZT12 (Figure 2), likely reflecting clearance of 8-oxo-dG during sleep. Our previous work demonstrated that increasing the time rats spent awake due to sleep disruption contributed to increases in 8-oxo-dG intensity, especially in PV-expressing neurons [48]. The mitochondria of PV-expressing neurons in particular exhibit elevated energy demands, as these neurons are essential for generating gamma activity, which requires a high and sustained firing rate and frequency [17], rendering PV cells especially vulnerable to oxidative stress compared with other types of neurons [110]. As demonstrated here, accumulation of oxidative stress occurred even without sleep disruption, but it is likely that both sleep and circadian mechanisms contribute to the rhythmicity of ROS accumulation and damage related to oxidative stress, as explained by the two-process model [111, 112]. Studying the dynamics of PNNs and PV neuronal activity from the framing of the two process-model may be helpful in understanding their variability, and examination of this is ongoing.

Both CLOCK and OTX2 appear to protect cells against the effects of oxidative stress [82, 89], and, as they regulate plasticity during development, they may also regulate diurnal changes. PNNs protect their underlying neurons from oxidative stress due to their enrichment in chondroitin sulfate [113], although PNNs are also subject to deleterious effects by oxidative stress [114]. Our findings that PNN intensity changes across the day/night cycle suggest that protective mechanisms from oxidative stress is also be subject to diurnal variation.

## Conclusions

Our findings indicate that PNN, PV, and 8-oxo-dG intensities fluctuate throughout the day. Increases in WFA and PV intensity precede changes in 8-oxo-dG intensity, suggesting a circadian-regulated increase in PNNs around PV cells at the onset of the active phase, potentially in preparation for increased glutamatergic input and presumed increases in cell firing. OTX2 may serve a role in coordinating diurnal changes in gene expression needed for optimal performance of PV cell firing during sleep and wakefulness, including proteins providing protection from oxidative stress. Diurnal fluctuation in these parameters are important and thus far are largely unconsidered factors in interpretation of the impact these fluctuations may have on basic cellular functions such as electrophysiological properties and transcription-coupling mechanisms typically measured during the light phase. Furthermore, interpretation of diurnal fluctuations in PV cell function is vital for considering treatment and study of disease states impacted by sleep and circadian factors. Given the fundamental role of PV-expressing neurons in brain disorders and the maintenance of cortical excitatory:inhibitory balance and gamma oscillations necessary for attention and cognitive flexibility [13, 17, 115], it is important to establish how daily rhythmicity is coordinated among OTX2, oxidative stress, and PNN/PV function in the mPFC.

## Acknowledgements

The authors would like to thank Dr. Megan Slaker, Ryan P. Todd, Monica Chang, and Nathan Allen for help with earlier stages of the experiments and Jonathan Ramos for assistance with analysis. This work was funded by Washington State University Alcohol and Drug Abuse Research Program and National Institutes of Health Grants DA 033404, DA 040965, and GM 134789.

## Ethics approval

All procedures were performed in accordance with the National Institutes of Health's Guidelines for the Care and Use of Laboratory Animals and with approval from the Institutional Animal Care and Use Committee at Legacy Research Institute and Washington State University.

## Availability of data and material

Data are available upon request.

## Code availability

Available for download at https://pipsqueak.ai

## Authors’ contributions

JHH and AEE, ETJ, and DH conducted experiments and analyzed data; PNB analyzed data; JPW performed statistical analyses and contributed to writing the manuscript; AD and AP contributed to writing the manuscript; and SAA, TEB, and BAS contributed to writing the manuscript, and designed and directed the experiments.

